# X-ray diffraction from hydrated macromolecular crystals using laser-driven X-ray source

**DOI:** 10.64898/2026.10.19.760121

**Authors:** Iuliia Baranova, Thomas Poulis, Roya Garayeva, Yelyzaveta Pulnova, Jaroslav Nejdl, Jakob Andreasson, Krishna Khakurel

## Abstract

Laser-driven X-ray sources offer a compact route towards laboratory-scale ultrafast structural dynamics. However, due to the low photon flux available from laboratory-scale sources, and the weak diffraction signal from biological specimens, their application to such samples remains challenging. Here, we demonstrate X-ray diffraction from Z-DNA crystals using a laser-driven Cu K*α* source (*λ* = 1.54 Å). Crystals were grown either directly in a custom-designed sample holder, which preserved the hydrated crystallisation environment and allowed *in situ* diffraction measurements, or using the hanging-drop vapour-diffusion method, followed by loop mounting for diffraction measurements. Well-defined Bragg reflections extending to approximately 2.5 Å resolution were recorded in 1–3 min exposures at a photon flux of approximately 2.2 × 10^5^ photons s^−1^ at the sample. The acquisition time and diffraction quality compare favourably with earlier demonstrations of macromolecular crystal diffraction using laser-driven X-ray sources. These results demonstrate the potential of modern compact laser-driven X-ray sources for structural studies of macromolecular crystals and provide a basis for their future application to laboratory-scale time-resolved crystallography.

**Synopsis:** Bragg diffraction to approximately 2.5 Å was recorded in 1–3 min exposures from hydrated Z-DNA crystals using a laser-driven Cu K*α* X-ray source, both for crystals grown *in situ* in a custom holder and for loop-mounted crystals. This is a step towards complete data collection and time-resolved crystallography of macromolecules with compact laser-driven sources.

## 1 Introduction

X-ray crystallography has played a central role in determining the three-dimensional structures of biological macromolecules at atomic resolution (Jaskolski *et al*., 2014). Conventional laboratory X-ray sources provide reliable access to structural measurements (Skarzynski, 2013), while synchrotron radiation and X-ray free-electron lasers (XFELs) provide high photon flux and, in the case of XFELs, ultrashort pulse durations that enable time-resolved experiments at picosecond and sub-picosecond timescales (Emma *et al*., 2010; Chapman *et al*., 2011). In particular, time-resolved crystallographic methods have made it possible to investigate structural changes in biological systems following optical or chemical excitation on timescales ranging from milliseconds to femtoseconds (Schotte *et al*., 2003; Tenboer *et al*., 2014; Barends *et al*., 2015). Access to large-scale X-ray facilities is, however, limited by strong competition for beamtime, motivating the development of compact sources capable of bringing ultrafast structural measurements into the laboratory.

Laser-driven X-ray sources provide a promising route towards this goal. When an intense ultrashort laser pulse is focused onto a solid target, the resulting laser-produced plasma emits hard X-rays, including characteristic emission lines suitable for diffraction experiments (Murnane *et al*., 1991; Rousse *et al*., 1994). The short timescale associated with laser-plasma X-ray generation additionally makes these sources attractive for pump–probe measurements of structural dynamics (Rousse *et al*., 2001; Reich *et al*., 2000). Furthermore, using the same laser source for X-ray generation and optical excitation reduces the timing jitter between pump and probe pulses (Elsaesser & Woerner, 2014). Laser-driven X-ray sources have been successfully employed in diffraction and imaging experiments on a range of solid-state systems (Rose-Petruck *et al*., 1999; Sokolowski-Tinten *et al*., 2003; Bargheer *et al*., 2006; Afshari *et al*., 2020; Pulnova *et al*., 2025). However, their application to macromolecular crystallography is considerably more demanding, because the moderate photon flux of compact plasma X-ray sources limits measurements of weakly scattering samples and places stringent requirements on the achievable signal-to-background ratio (Bonvalet *et al*., 2006; Baranova *et al*., 2026; Khakurel *et al*., 2024*b*). Macromolecular crystals therefore represent a particularly challenging class of samples for such sources. Early experiments demonstrated that diffraction from protein crystals could be obtained using laser-produced X-ray sources, establishing the feasibility of laboratory-scale ultrafast macromolecular crystallography (Bonvalet *et al*., 2006). Nevertheless, the limited number of photons available at the sample resulted in relatively long acquisition times and restricted diffraction quality. Recent advances in high-power laser technology, X-ray generation and collection efficiency, source stability, repetition rate, and detector sensitivity have improved the prospects for measurements of weakly scattering samples with compact laser-driven sources (Baranova *et al*., 2026), motivating their further application to biological crystallography.

Z-DNA provides an attractive model system for evaluating such sources. In contrast to the canonical right-handed B-DNA double helix, Z-DNA adopts a left-handed helical conformation characterised by a zig-zag arrangement of the phosphodiester backbone (Rich & Zhang, 2003). The first atomic structure of Z-DNA was determined from crystals of the self-complementary DNA hexamer d(CGCGCG), establishing the structural basis of this unusual DNA conformation (Wang *et al*., 1979). Since then, the d(CGCGCG)_2_ duplex has become a well-characterised model system in nucleic-acid crystallography, with structures determined at exceptionally high resolution (Gessner *et al*., 1989; Brzezinski *et al*., 2011). Its small and well-established unit cell, together with the extensive structural information available for comparison, makes crystalline Z-DNA particularly suitable as a benchmark biological system for evaluating emerging X-ray sources. Previous studies of ultra-fast dynamics in nucleotides report dynamics on timescales ranging from a few picoseconds to tens of picoseconds (Crespo-Hernández *et al*., 2004; Hare *et al*., 2007; Middleton *et al*., 2009; Prokhorenko *et al*., 2016; Kufner *et al*., 2020). Z-DNA is therefore also a potential model system for extending the present work to transient structural dynamics.

A further challenge in macromolecular crystallography is the preservation of crystal integrity during sample preparation and mounting. Macromolecular crystals contain substantial amounts of solvent (Matthews, 1968) and can be sensitive to dehydration and mechanical manipulation during harvesting and mounting (Fischer, 2021). Growing crystals directly within an X-ray-compatible sample environment provides an alternative strategy that minimises sample manipulation and allows the crystal to remain in its hydrated crystallisation conditions throughout the experiment (le Maire *et al*., 2011; Axford *et al*., 2012).

In this work, we demonstrate diffraction from Z-DNA crystals using a laser-driven X-ray source producing Cu K*α* radiation at a wavelength of 1.54 Å. The experiment builds on the recently developed laser-driven diffraction platform at ELI Beamlines (Pulnova *et al*., 2025; Baranova *et al*., 2026). For *in situ* experiments, the crystals were grown and measured directly within a custom sample holder designed to preserve their hydrated crystallisation environment and avoid crystal harvesting, following the general advantages of *in situ* macromolecular crystallography (Axford *et al*., 2012; Fischer, 2021). Diffraction patterns were recorded using a JUNGFRAU pixel detector (Leonarski *et al*., 2018) and compared with measurements obtained using a conventional X-ray tube source. Multiple well-defined Bragg reflections were observed, extending to approximately 2.5 Å resolution. These measurements demonstrate the capability of a compact laser-driven X-ray source to access crystallographically relevant length scales in a biological crystal and represent a step towards laboratory-scale time-resolved structural studies of biological macromolecules.

## 2 Experiment

### 2.1 Experimental setup

The diffraction measurements were performed using the laser-driven X-ray source at the TREX end station of the ELI Beamlines facility (Fig. 1), which produces characteristic Cu K*α* radiation at a wavelength of *λ* = 1.54 Å (*E ≈* 8.05 keV) (Pulnova *et al*., 2025). The femtosecond laser system delivered pulses with an energy of approximately 10 mJ, a central wavelength of 800 nm, and a duration of 45 fs at a repetition rate of 1 kHz. The laser pulses were focused to a spot of approximately 8 *µ*m on a copper tape target, generating a laser-produced plasma with an X-ray source size of approximately 10 *µ*m. The emitted Cu K*α* radiation was spectrally filtered and focused onto the sample by a multilayer Montel optic. The X-ray beam size at the sample position was approximately 100 *µ*m (FWHM) in both directions, and the estimated photon flux at the sample was approximately 2.2 *×* 10^5^ photons s^*−*1^.

**Figure 1:**
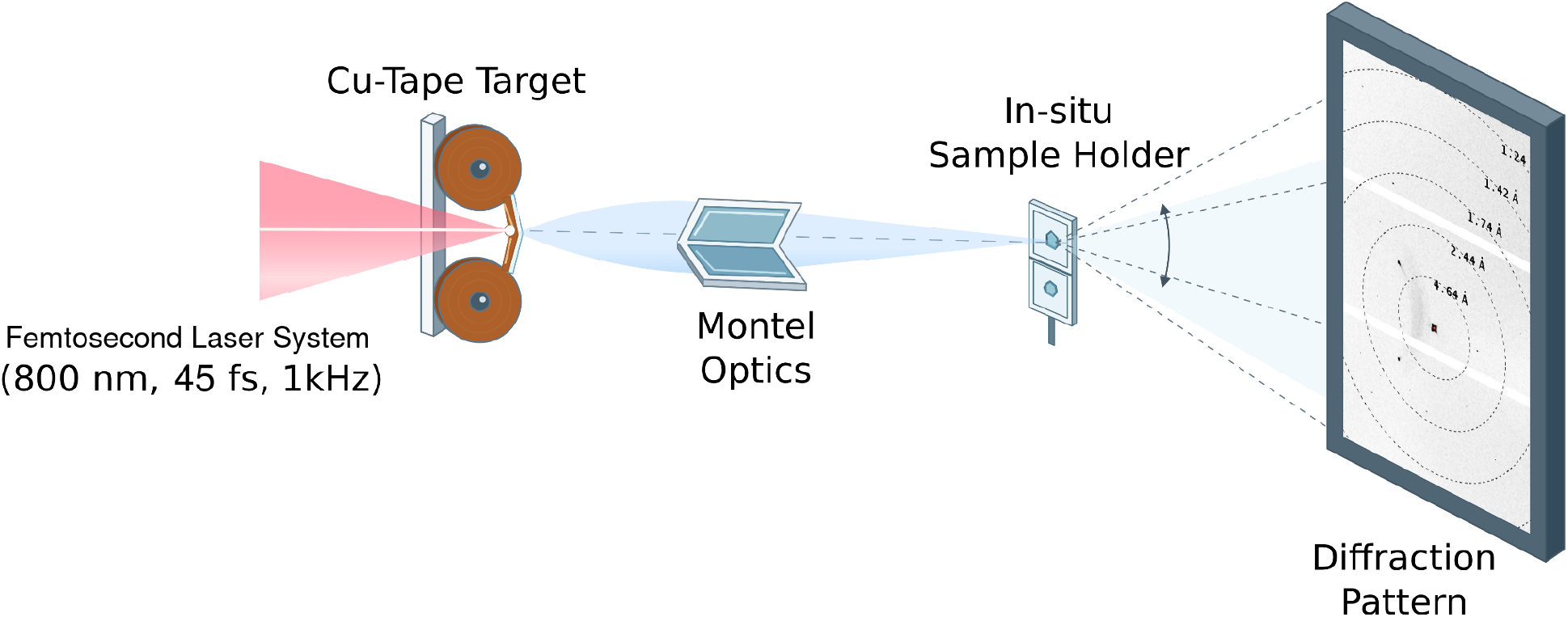
Schematic of the experimental setup. Pulses from the femtosecond laser system are focused onto a copper tape target, generating Cu K*α* radiation, which is collected and focused onto the sample by a multilayer Montel optic. The diffraction pattern from the crystal in the *in situ* sample holder is recorded by the detector.

For the measurements presented here, each diffraction pattern was accumulated over an exposure time of 1–3 min, corresponding to approximately 6 *×* 10^4^–1.8 *×* 10^5^ laser shots.

### 2.2 Z-DNA crystallisation and *in situ* sample environment

For *in situ* experiments, Z-DNA crystals were grown directly within a custom-designed sample holder that served simultaneously as the crystallisation well and the diffraction mount. The custom sample holder was fabricated using 3D printing from a polymer material and equipped with X-ray-transparent windows made of 20 *µ*m-thick cyclic olefin copolymer (COC).

The self-complementary DNA oligonucleotide d(CGCGCG) was crystallised following a previously described protocol (Khakurel *et al*., 2024*a*). Lyophilised DNA hexamer was dissolved in Milli-Q water to a concentration of 50 mg/mL. The crystallisation buffer was prepared using MES [2-(N-morpholino)ethanesulfonic acid], ammonium sulfate, and magnesium acetate at final concentrations of 50 mM, 2.5 M and 10 mM, respectively, with a final pH of 6.1. The DNA solution and crystallisation buffer were mixed at a 1:1 volume ratio, giving a final drop volume of 1 *µ*L in the sample holder.

In addition to the *in situ* sample holders, crystals were grown using the hanging-drop vapour-diffusion method (Benvenuti & Mangani, 2007) and subsequently mounted in loops, as is commonly done in single-crystal X-ray diffraction experiments. To minimise dehydration, the loop-mounted crystal was enclosed in a polyester capillary containing mother liquor. All measurements were performed at room temperature. Diffraction from Z-DNA crystals was obtained with both approaches.

Representative optical micrographs of crystals grown within the sample holder and of a loop-mounted crystal are shown in Figs. 2(a) and 2(f), respectively. The crystal used for the diffraction measurements had approximate dimensions of 200 *µ*m *×* 200 *µ*m. In a separate experiment under the same crystallisation conditions, the crystals were found to belong to the orthorhombic space group *P* 2_1_2_1_2_1_, consistent with previous reports (Wang *et al*., 1979; Brzezinski *et al*., 2011).

**Figure 2:**
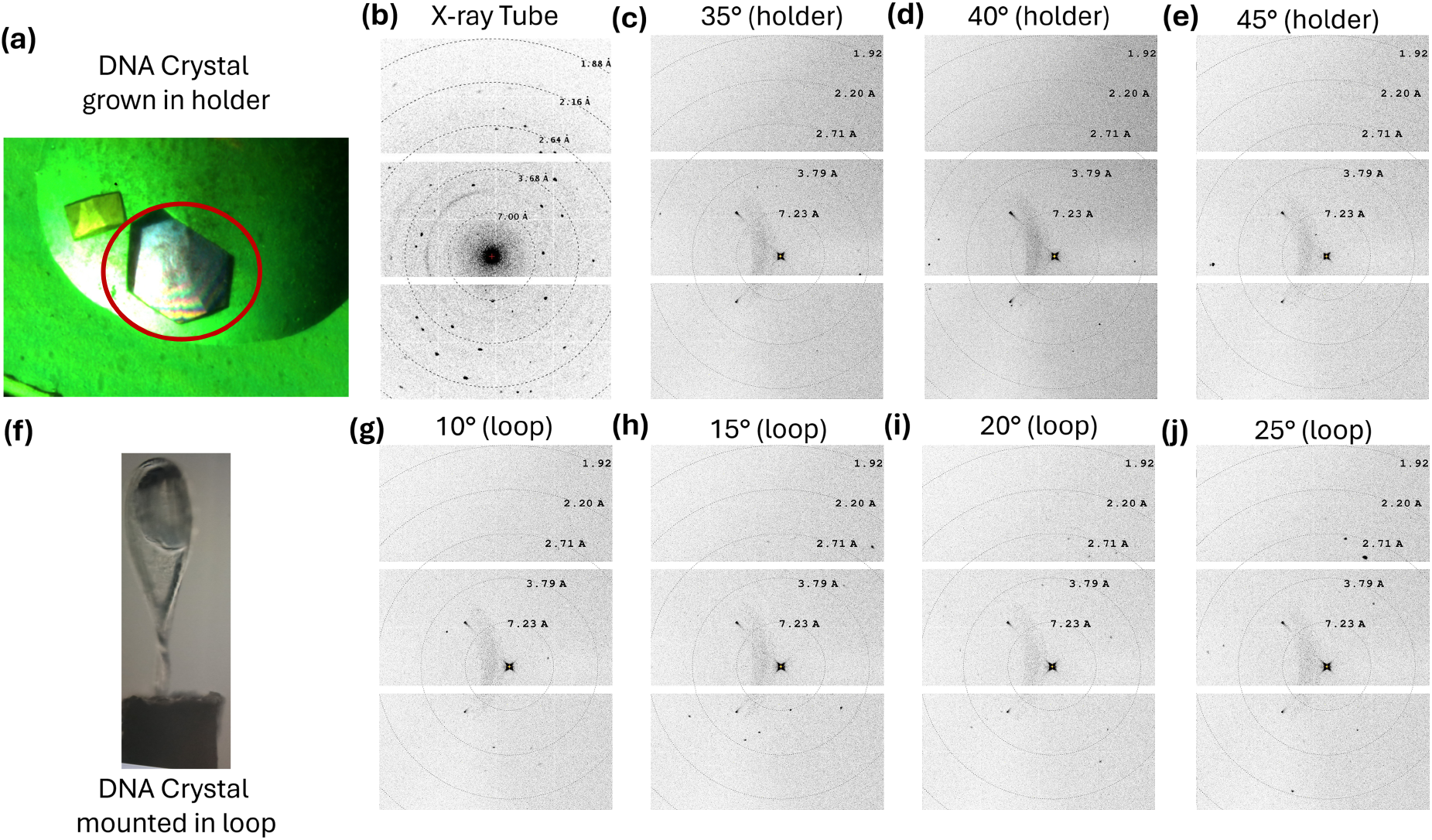
(a) Optical image of Z-DNA crystals grown in the *in situ* sample holder. The crystal selected for X-ray diffraction measurements is marked with a red circle. (b) Diffraction pattern of Z-DNA recorded with the conventional X-ray tube. (c)–(e) Diffraction patterns of a Z-DNA crystal grown in the *in situ* holder, recorded with the laser-driven X-ray source at different orientations. (f) Optical image of a Z-DNA crystal mounted in a loop. (g)–(j) Diffraction patterns of the loop-mounted Z-DNA crystal recorded with the laser-driven X-ray source at different orientations. Angles are relative rotation angles with an arbitrary zero. Dotted circles indicate resolution rings calculated from the LaB_6_-calibrated geometry. Each pattern was accumulated over 1–3 min.

### 2.3 X-ray diffraction measurements

X-ray diffraction patterns were recorded using a JUNGFRAU detector (Mozzanica *et al*., 2018; Leonarski *et al*., 2018) consisting of three modules (approximately 1.5 million pixels, 75 *µ*m pixel size), positioned at a sample-to-detector distance of 67.7 mm. The detector was operated with an integration time of 1 *µ*s per frame, using the gain settings for 8 keV photons. The detector distance was calibrated using the diffraction pattern of a lanthanum hexaboride (LaB_6_) standard, and the resolution of each reflection was obtained from its position on the detector using this calibrated geometry and Bragg’s law. The crystal was kept at a fixed orientation during each acquisition, so that each measurement is a still diffraction pattern sampling a limited region of reciprocal space. Patterns were recorded at several crystal orientations; the angles given in Figs. 2 and 3 are relative rotation angles with an arbitrary zero.

**Figure 3:**
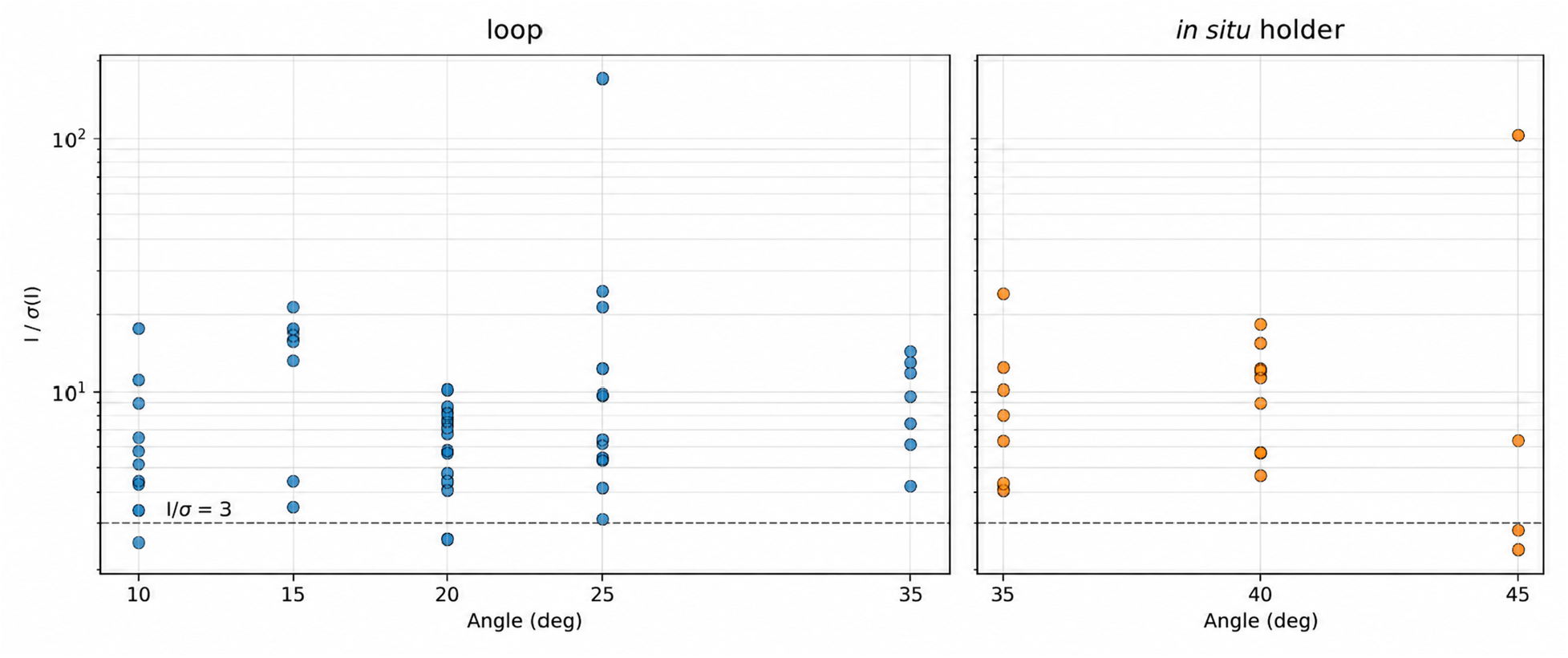
*I/σ*(*I*) of peaks extracted from laser-driven diffraction patterns recorded from the crystal mounted in a loop (left) and from the crystal grown in the *in situ* holder (right), plotted against the relative crystal rotation angle (arbitrary zero). The dashed line marks *I/σ*(*I*) = 3; 70 of the 74 peaks lie above it.

Diffraction patterns were recorded with both the laser-driven X-ray source and a conventional X-ray tube. Both sources were equipped with identical multilayer Montel optics, providing comparable X-ray beam parameters at the sample position. The conventional-source measurements served as a reference for assessing crystal quality and for identifying the crystallographic origin of the reflections observed under laser-driven illumination. For the conventional source, the photon flux at the sample was approximately seven times higher than for the laser-driven source.

The diffraction patterns were analysed using the Circle Plot tool in ALBULA (Dectris). Individual Bragg reflections were selected, and their integrated intensities were determined within circular regions centred on the reflections. The local background was estimated from the surrounding region and subtracted from the integrated signal. The background-corrected integrated intensities, *I*, and their uncertainties, *σ*(*I*), were obtained directly from ALBULA and used to calculate the signal-to-noise ratio, *I/σ*(*I*). Additionally, one-dimensional intensity profiles of representative reflections were extracted and fitted with Gaussian functions using OriginPro. For these profiles, the uncertainties were estimated assuming Poisson counting statistics, as the square root of the measured photon counts. Both the intensities and their uncertainties were subsequently normalised to the acquisition time and expressed in photons per second (ph/s).

## 3 Results and Discussion

### 3.1 X-ray diffraction from Z-DNA

Crystals grown in the custom holder remained in their hydrated crystallisation environment throughout the experiment, so no harvesting or transfer was needed before the measurement. For comparison, crystals grown by hanging-drop vapour diffusion were mounted in a loop and enclosed in a polyester capillary containing mother liquor, which also kept them hydrated.

A representative diffraction pattern recorded with the conventional X-ray tube in a 1-min exposure is shown in Fig. 2(b). Multiple discrete Bragg reflections are clearly visible above the background, confirming the quality of the crystal. After this confirmation, diffraction patterns were recorded with the laser-driven X-ray source. Representative patterns at different crystal orientations are shown in Fig. 2: the first row [Figs. 2(c)–2(e)] shows patterns from a Z-DNA crystal grown in the *in situ* holder, and the second row [Figs. 2(g)–2(j)] shows patterns from a loop-mounted crystal grown using the conventional hanging-drop vapour-diffusion method. In both cases, the reflections are spatially localised and exhibit the spot-like profiles expected for diffraction from a single crystal.

The highest-resolution reflections observed in the present measurements correspond to *d*-spacings of approximately 2.5 Å, as determined from the LaB_6_-calibrated detector geometry. This resolution is encouraging given that the measurements were performed with a compact laser-driven source and a biological crystal kept in its crystallisation environment. Although the present experiment does not constitute a complete crystallographic dataset, the observation of well-defined reflections at approximately 2.5 Å shows that the source provides sufficient X-ray flux and signal-to-background ratio to access crystallographically relevant information from the Z-DNA crystal.

### 3.2 Quantitative characterisation of the diffraction signal

To characterise the laser-driven diffraction beyond visual inspection of the detector images, individual Bragg reflections were analysed in terms of their integrated intensities, signal-to-noise ratios, and spatial widths. Fig. 3 shows *I/σ*(*I*) for peaks extracted from patterns recorded from the loop-mounted crystal (left) and from the crystal grown in the *in situ* holder (right). A detailed statistical analysis of the peaks is given in the supporting information. Of the 74 reflections analysed (22 from the crystal in the holder and 52 from the loop-mounted crystal), 70 (95%) have *I/σ*(*I*) *>* 3, i.e., they are clearly distinguishable from the background noise (Borek *et al*., 2003).

The spatial profiles of the Bragg reflections were further characterised by fitting Gaussian functions with a constant background (Fig. 4). The fitted width provides a quantitative measure of the spot size on the detector. Most of the representative profiles, recorded with the crystal in the loop and in the *in situ* holder, are well described by Gaussian functions, and the widths obtained with the laser-driven source are comparable to those obtained with the X-ray tube. Because the spot width in a still image is largely determined by the beam convergence and the illuminated crystal volume (Nave, 1998), these widths characterise spot quality rather than crystal mosaicity.

**Figure 4:**
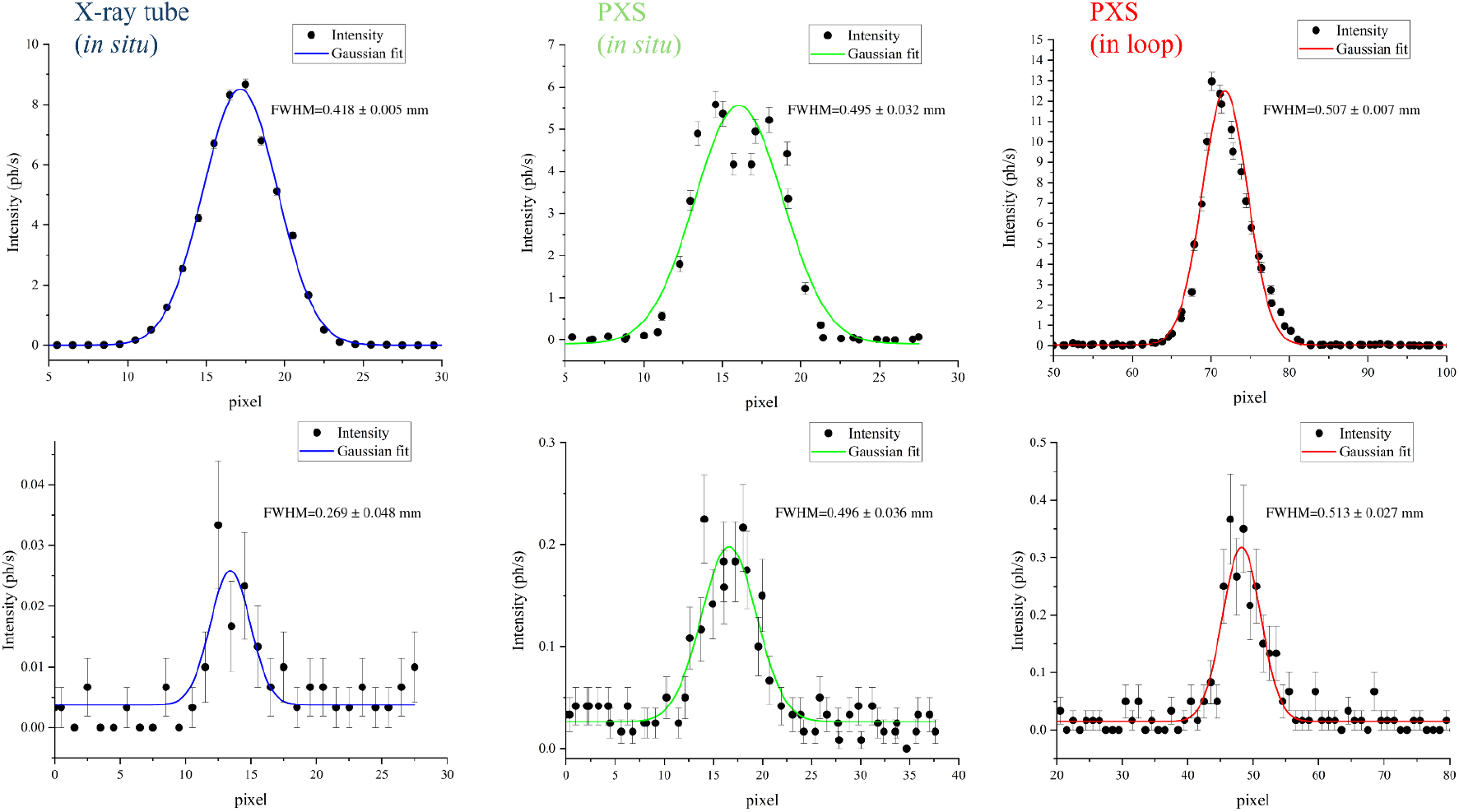
Intensity profiles of representative Bragg reflections (black circles) with Gaussian fits (solid lines), recorded using a conventional X-ray tube with a crystal grown *in situ* in the holder (left column), a laser-driven plasma X-ray source (PXS) with a crystal grown *in situ* in the holder (middle column), and PXS with a loop-mounted crystal (right column). The top row shows the intensity profiles of the strongest reflections, and the bottom row shows the intensity profiles of the weakest reflections. Error bars represent uncertainties in the measured intensities. Horizontal axes are in detector pixels (1 pixel = 75 *µ*m). The full width at half maximum (FWHM) values obtained from the Gaussian fits are indicated in each panel in millimetres (mm).

Together, the integrated intensities, *I/σ*(*I*) values, and spot widths provide complementary measures of diffraction quality. The values obtained from the laser-driven data indicate that collecting complete datasets with this source would allow further crystallographic analysis.

### 3.3 Implications for laboratory-scale time-resolved macromolecular crystallography

The present measurements demonstrate that a state-of-the-art laser-driven Cu K*α* source can produce diffraction to approximately 2.5 Å from Z-DNA crystals, whether grown *in situ* in the custom holder or by hanging-drop vapour diffusion and then loop-mounted. This extends earlier demonstrations of diffraction from biological samples and protein crystals with laser-produced X-ray sources (Frankel & Forsyth, 1979; Bonvalet *et al*., 2006) and represents an important step towards the use of compact laser-driven X-ray sources for structural measurements of biological samples.

The *in situ* sample environment is an important component of the experimental approach. Biological crystals are sensitive to changes in hydration and to mechanical manipulation during harvesting and mounting (Axford *et al*., 2012; Fischer, 2021). Growing the crystal directly within the diffraction holder minimises these handling steps and maintains the crystal in its hydrated crystallisation environment during the measurement. The approach also provides a reproducible geometry that may be advantageous for future experiments involving repeated or time-resolved measurements.

Extending the current experiment to sample a larger region of reciprocal space would enable indexing and, ultimately, structure determination. Given the known *P* 2_1_2_1_2_1_ symmetry and relatively small unit cell of the Z-DNA crystal (Wang *et al*., 1979; Brzezinski *et al*., 2011), acquiring indexed diffraction patterns at a series of orientations provides a plausible route towards a more complete crystallographic dataset (White *et al*., 2012; Winter *et al*., 2018).

The broader motivation for developing laser-driven X-ray sources for biological crystallography is their potential for time-resolved structural measurements (Rousse *et al*., 2001; Bonvalet *et al*., 2006; Afshari *et al*., 2020). Because X-ray generation is intrinsically synchronised to the driving laser pulse, such sources are naturally suited to optical-pump/X-ray-probe experiments (Rousse *et al*., 2001; Afshari *et al*., 2020). Further improvements in photon flux, source stability, acquisition rate, sample positioning, and reciprocal-space sampling will be required before complete time-resolved crystallographic datasets can be routinely obtained. Nevertheless, the observation of diffraction to approximately 2.5 Å from Z-DNA shows that compact laser-driven sources are approaching a regime in which structurally relevant measurements of biological crystals can be performed in the laboratory.

## 4 Conclusions

We have demonstrated X-ray diffraction from Z-DNA crystals using a laser-driven Cu K*α* source operating at a wavelength of 1.54 Å. A custom sample holder allowed the crystal to be grown and measured *in situ* within its hydrated crystallisation environment, eliminating the need for crystal harvesting and transfer prior to the diffraction experiment.

The laser-driven measurements reveal multiple well-defined Bragg reflections extending to approximately 2.5 Å resolution. Comparison with measurements performed using a conventional X-ray tube source supports the crystallographic origin of the observed reflections. The analysis of reflection intensities, *I/σ*(*I*) values, and peak widths indicates that the diffraction quality is sufficient for further crystallographic analysis once more complete datasets are collected.

The present experiment was performed in a still-image geometry and therefore samples only a limited region of reciprocal space. Since the known crystallographic parameters of the Z-DNA crystal provide a basis for simulating and indexing the observed diffraction patterns, measurements at multiple crystal orientations would substantially increase reciprocal-space coverage and provide a pathway towards the collection of more complete diffraction datasets. We will pursue these experiments as the next step of this research.

Taken together, these results demonstrate the increasing potential of compact laser-driven X-ray sources for structural studies of biological crystals. Further improvements in source brightness, acquisition efficiency, and reciprocal-space sampling, combined with the intrinsically short timescale of laser-driven X-ray generation, may ultimately enable laboratory-scale time-resolved studies of the structural dynamics of biological macromolecules. The use of mid-infrared driving lasers for efficient X-ray generation (Weisshaupt *et al*., 2014; Koç *et al*., 2021), together with the development of high-repetition-rate lasers, shows promise for meeting the photon flux requirements of such experiments.

## Acknowledgements

The experiments presented in this article were performed as part of ELI user proposal eli50124.

## Funding

This work was partially funded by ELI ERIC.

### Data availability

The data supporting the results reported in this article are available from the corresponding author upon reasonable request.

