## Supplemental Information for "X-ray diffraction from hydrated macromolecular crystals using laser-driven X-ray source"

### Supporting information for: X-ray diffraction from hydrated macromolecular crystals using laser-driven X-ray source

#### S1 Integrated intensities of individual Bragg reflections

Tables S2–S10 list all Bragg reflections analysed in the still diffraction patterns shown in Figs. 2 and 3 of the main text. Integrated intensities were determined with the Circle Plot tool in ALBULA (Dectris). Each reflection was enclosed in a circular integration region, the mean background per pixel was estimated from the surrounding region and subtracted, and the background-corrected integrated intensity $I$ and its uncertainty $\sigma(I)$ were obtained as described in Section 2.3 of the main text. Intensities are given in detector counts accumulated over the full exposure, without normalisation to the acquisition time.

Rotation angles are relative, with an arbitrary zero. The angles for the crystal in the *in situ* holder and for the loop-mounted crystal refer to different crystals and cannot be compared with each other. Reflection numbers are labels within each pattern only.

Table S1 summarises the data. Of the 74 reflections measured with the laser-driven X-ray source, 70 (95%) have $I/\sigma(I) > 3$.

Table S1: Summary of the analysed diffraction patterns. PXS: laser-driven plasma X-ray source. $N$: number of reflections analysed. The column definitions for Tables S2–S10 are given below the table.

| Table | Source | Crystal | Angle (°) | Exposure (min) | $N$ | $N$ with $I/\sigma(I) > 3$ | Median $I/\sigma(I)$ | Max. $I/\sigma(I)$ |
|---|---|---|---|---|---|---|---|---|
| S2 | PXS | Holder | 35 | 3 | 8 | 8 | 7.3 | 24.4 |
| S3 | PXS | Holder | 40 | 2 | 10 | 10 | 11.5 | 18.3 |
| S4 | PXS | Holder | 45 | 1 | 4 | 2 | 4.6 | 102.8 |
| S5 | PXS | Loop | 10 | 1 | 10 | 9 | 5.5 | 17.8 |
| S6 | PXS | Loop | 15 | 1 | 8 | 8 | 15.8 | 21.5 |
| S7 | PXS | Loop | 20 | 1 | 15 | 14 | 7.3 | 10.2 |
| S8 | PXS | Loop | 25 | 1 | 12 | 12 | 8.1 | 171.4 |
| S9 | PXS | Loop | 35 | 1 | 7 | 7 | 9.6 | 14.4 |
| All PXS patterns | | | | | 74 | 70 | 7.6 | 171.4 |
| S10 | X-ray tube | Holder | 10 | 1 | 2[a] | 2 | – | 302.3 |

[a] Only reflections No. 1 and No. 25 of this pattern are listed.

Column definitions for Tables S2–S10: $I$, background-corrected integrated intensity (counts); $\sigma(I)$, uncertainty of $I$ (counts); BG, mean background per pixel (counts); Mean, mean value per pixel within the integration region (counts); Area, integration area (pixels; 1 pixel = $75 \times 75$ µm$^2$); Max., highest pixel value within the integration region (counts).

Table S2: PXS, crystal grown in the *in situ* holder, rotation angle 35°, exposure 3 min. Image file `pos1_35deg_3min_3`.

| No. | $I$ | $\sigma(I)$ | $I/\sigma(I)$ | BG | Mean | Area | Max. |
|---|---|---|---|---|---|---|---|
| 1 | 421.75 | 33.77 | 12.49 | 1.55 | 3.65 | 210.06 | 18 |
| 2 | 475.69 | 46.65 | 10.20 | 2.45 | 4.32 | 254.47 | 19 |
| 3 | 216.32 | 33.86 | 6.39 | 2.41 | 3.82 | 153.94 | 13 |
| 4 | 148.77 | 36.41 | 4.09 | 2.67 | 3.64 | 153.94 | 12 |
| 5 | 130.11 | 31.93 | 4.07 | 2.67 | 3.52 | 153.94 | 12 |
| 6 | 1230.23 | 50.48 | 24.37 | 3.02 | 10.33 | 201.06 | 42 |
| 7 | 175.73 | 40.58 | 4.33 | 2.90 | 3.78 | 201.06 | 14 |
| 8 | 214.24 | 26.34 | 8.13 | 1.31 | 2.70 | 153.94 | 12 |

Table S3: PXS, crystal grown in the *in situ* holder, rotation angle 40°, exposure 2 min. Image file `pos1_40deg_2min_5`.

| No. | $I$ | $\sigma(I)$ | $I/\sigma(I)$ | BG | Mean | Area | Max. |
|---|---|---|---|---|---|---|---|
| 1 | 459.90 | 39.17 | 11.74 | 2.11 | 4.41 | 201.06 | 19 |
| 2 | 822.94 | 44.98 | 18.29 | 1.95 | 5.19 | 254.47 | 31 |
| 3 | 606.28 | 39.18 | 15.48 | 2.04 | 5.06 | 201.06 | 19 |
| 4 | 181.91 | 31.95 | 5.69 | 2.58 | 3.76 | 153.94 | 16 |
| 5 | 631.51 | 51.73 | 12.21 | 3.62 | 6.11 | 254.47 | 27 |
| 6 | 593.73 | 52.75 | 11.25 | 4.83 | 7.79 | 201.06 | 28 |
| 7 | 346.11 | 38.55 | 8.98 | 3.80 | 6.05 | 153.94 | 15 |
| 8 | 501.89 | 41.69 | 12.04 | 3.23 | 6.50 | 153.94 | 24 |
| 9 | 134.85 | 28.91 | 4.66 | 3.52 | 5.25 | 78.54 | 15 |
| 10 | 140.66 | 24.60 | 5.72 | 3.47 | 6.29 | 50.27 | 18 |

Table S4: PXS, crystal grown in the *in situ* holder, rotation angle 45°, exposure 1 min. Image file `pos1_45deg_1min_7`.

| No. | $I$ | $\sigma(I)$ | $I/\sigma(I)$ | BG | Mean | Area | Max. |
|---|---|---|---|---|---|---|---|
| 1 | 199.63 | 31.16 | 6.41 | 2.12 | 3.89 | 113.10 | 14 |
| 2 | 11516.50 | 111.97 | 102.85 | 1.88 | 59.29 | 201.06 | 346 |
| 3 | 63.79 | 26.86 | 2.37 | 1.70 | 2.23 | 113.10 | 10 |
| 4 | 71.18 | 25.20 | 2.82 | 1.74 | 2.20 | 153.94 | 10 |

Table S5: PXS, loop-mounted crystal, rotation angle 10°, exposure 1 min. Image file `10deg_1min_50`.

| No. | $I$ | $\sigma(I)$ | $I/\sigma(I)$ | BG | Mean | Area | Max. |
|---|---|---|---|---|---|---|---|
| 1 | 605.69 | 34.12 | 17.75 | 1.13 | 4.15 | 201.06 | 28 |
| 2 | 247.94 | 37.58 | 6.60 | 1.88 | 2.85 | 254.47 | 11 |
| 3 | 178.70 | 34.49 | 5.18 | 2.07 | 2.96 | 201.06 | 9 |
| 4 | 323.08 | 29.19 | 11.07 | 1.46 | 3.57 | 153.94 | 16 |
| 5 | 298.92 | 33.32 | 8.97 | 1.31 | 2.49 | 254.47 | 12 |
| 6 | 221.61 | 37.99 | 5.83 | 1.54 | 2.24 | 314.16 | 11 |
| 7 | 163.71 | 37.96 | 4.31 | 1.79 | 2.44 | 254.47 | 10 |
| 8 | 87.93 | 26.15 | 3.36 | 1.22 | 1.66 | 201.06 | 11 |
| 9 | 108.91 | 24.42 | 4.46 | 1.26 | 1.97 | 153.94 | 8 |
| 10 | 51.58 | 20.33 | 2.54 | 1.07 | 1.41 | 153.94 | 6 |

Table S6: PXS, loop-mounted crystal, rotation angle 15°, exposure 1 min. Image file `15deg_1min_43`.

| No. | $I$ | $\sigma(I)$ | $I/\sigma(I)$ | BG | Mean | Area | Max. |
|---|---|---|---|---|---|---|---|
| 1 | 1010.94 | 63.12 | 16.02 | 2.43 | 4.33 | 530.93 | 26 |
| 2 | 138.02 | 39.36 | 3.51 | 2.29 | 2.83 | 254.47 | 12 |
| 3 | 763.91 | 42.87 | 17.82 | 1.89 | 4.90 | 254.47 | 25 |
| 4 | 594.34 | 34.89 | 17.03 | 1.34 | 4.30 | 201.06 | 21 |
| 5 | 807.65 | 37.58 | 21.49 | 1.24 | 5.27 | 201.06 | 32 |
| 6 | 456.78 | 34.78 | 13.13 | 1.09 | 2.54 | 314.16 | 23 |
| 7 | 480.57 | 30.79 | 15.61 | 0.90 | 2.79 | 254.47 | 14 |
| 8 | 99.28 | 22.58 | 4.40 | 1.11 | 1.76 | 153.94 | 14 |

Table S7: PXS, loop-mounted crystal, rotation angle 20°, exposure 1 min. Image file `20deg_1min_37`.

| No. | $I$ | $\sigma(I)$ | $I/\sigma(I)$ | BG | Mean | Area | Max. |
|---|---|---|---|---|---|---|---|
| 1 | 315.40 | 37.27 | 8.46 | 1.48 | 2.49 | 314.16 | 12 |
| 2 | 320.86 | 36.46 | 8.80 | 1.74 | 3.00 | 254.47 | 12 |
| 3 | 192.02 | 33.64 | 5.71 | 1.90 | 2.86 | 201.06 | 13 |
| 4 | 151.51 | 31.90 | 4.75 | 1.49 | 2.08 | 254.47 | 8 |
| 5 | 311.35 | 41.39 | 7.52 | 1.49 | 2.31 | 380.13 | 13 |
| 6 | 247.50 | 30.71 | 8.06 | 1.46 | 3.20 | 201.06 | 16 |
| 7 | 340.47 | 33.52 | 10.16 | 1.73 | 3.42 | 201.06 | 13 |
| 8 | 279.59 | 27.36 | 10.22 | 0.82 | 1.92 | 254.47 | 9 |
| 9 | 116.57 | 16.98 | 6.87 | 0.71 | 1.75 | 113.10 | 8 |
| 10 | 146.13 | 18.87 | 7.74 | 0.81 | 2.11 | 113.10 | 9 |
| 11 | 118.92 | 20.26 | 5.87 | 0.86 | 1.63 | 153.94 | 13 |
| 12 | 49.67 | 18.93 | 2.62 | 1.21 | 1.65 | 113.10 | 13 |
| 13 | 176.76 | 24.31 | 7.27 | 0.83 | 1.71 | 201.06 | 11 |
| 14 | 105.70 | 24.04 | 4.40 | 0.85 | 1.38 | 201.06 | 9 |
| 15 | 98.16 | 23.93 | 4.10 | 1.16 | 1.65 | 201.06 | 8 |

Table S8: PXS, loop-mounted crystal, rotation angle 25°, exposure 1 min. Image file `25deg_1min_31`.

| No. | $I$ | $\sigma(I)$ | $I/\sigma(I)$ | BG | Mean | Area | Max. |
|---|---|---|---|---|---|---|---|
| 1 | 32391.00 | 188.92 | 171.45 | 2.08 | 47.93 | 706.86 | 778 |
| 2 | 1196.32 | 48.00 | 24.92 | 1615.13 | 8.05 | 201.06 | 37 |
| 3 | 396.71 | 32.41 | 12.24 | 1.74 | 3.72 | 201.06 | 14 |
| 4 | 294.21 | 30.47 | 9.65 | 1.64 | 3.56 | 153.94 | 14 |
| 5 | 760.23 | 35.36 | 21.50 | 1.06 | 4.85 | 201.06 | 25 |
| 6 | 164.75 | 25.50 | 6.46 | 1.52 | 2.98 | 113.10 | 10 |
| 7 | 277.87 | 28.10 | 9.89 | 0.95 | 2.34 | 201.06 | 11 |
| 8 | 71.05 | 22.82 | 3.11 | 1.08 | 1.71 | 113.10 | 7 |
| 9 | 145.79 | 23.64 | 6.17 | 0.93 | 1.65 | 201.06 | 7 |
| 10 | 136.71 | 25.72 | 5.31 | 1.19 | 1.87 | 201.06 | 8 |
| 11 | 130.43 | 31.33 | 4.16 | 1.81 | 2.46 | 201.06 | 13 |
| 12 | 114.69 | 20.93 | 5.48 | 1.51 | 2.53 | 113.10 | 7 |

Table S9: PXS, loop-mounted crystal, rotation angle 35°, exposure 1 min. Image file `35deg_1min_19`.

| No. | $I$ | $\sigma(I)$ | $I/\sigma(I)$ | BG | Mean | Area | Max. |
|---|---|---|---|---|---|---|---|
| 1 | 573.67 | 44.33 | 12.94 | 2.04 | 3.87 | 314.16 | 22 |
| 2 | 401.00 | 27.81 | 14.42 | 1.00 | 3.00 | 201.06 | 15 |
| 3 | 125.13 | 29.63 | 4.22 | 1.08 | 1.57 | 254.47 | 9 |
| 4 | 284.90 | 29.73 | 9.58 | 1.26 | 2.68 | 201.06 | 12 |
| 5 | 339.49 | 28.86 | 11.76 | 1.34 | 3.55 | 153.94 | 14 |
| 6 | 214.98 | 28.60 | 7.52 | 1.20 | 2.27 | 201.06 | 11 |
| 7 | 168.52 | 27.22 | 6.19 | 1.08 | 1.75 | 254.47 | 8 |

Table S10: Conventional X-ray tube, crystal grown in the *in situ* holder, rotation angle 10°, exposure 1 min. Only reflections No. 1 and No. 25 are listed. Image file `10deg_1min_50`.

| No. | $I$ | $\sigma(I)$ | $I/\sigma(I)$ | BG | Mean | Area | Max. |
|---|---|---|---|---|---|---|---|
| 1 | 92690.00 | 306.60 | 302.32 | 2.00 | 297.50 | 314.20 | 2680 |
| 25 | 53.73 | 14.56 | 3.69 | 1.57 | 2.65 | 50.27 | 10 |
